# Capturing cognitive events embedded in the real-world using mobile EEG and Eye-Tracking

**DOI:** 10.1101/2021.11.30.470560

**Authors:** Simon Ladouce, Magda Mustile, Frédéric Dehais

## Abstract

The study of cognitive processes underlying natural behaviours implies to depart from computerized paradigms and artificial experimental probes. The aim of the present study is to assess the feasibility of capturing neural markers of visual attention (P300 Event-Related Potentials) in response to objects embedded in a real-world environment. To this end, electroencephalography and eye-tracking data were recorded while participants attended stimuli presented on a tablet and while they searched for books in a library. Initial analyses of the library data revealed P300-like features shifted in time. A Dynamic Time Warping analysis confirmed the presence of P300 ERP in the library condition. Library data were then lag-corrected based on cross-correlation coefficients. Together these approaches uncovered P300 ERP responses in the library recordings. These findings highlight the relevance of scalable experimental designs, joint brain and body recordings and template-matching analyses to capture cognitive events during natural behaviours.

## 1 INTRODUCTION

The emergence of mobile brain and body imaging (MoBI, Gramann et al. (2011)) research methods provides the unprecedented opportunity to depart from artificial laboratory-based settings to study cognitive processes directly in real-world environments (Makeig et al., 2009; Gramann et al., 2014; De Vos et al., 2014). Over the last decade, technical advances have been made toward the miniaturization of sensors improving the portability of research-grade hardware (Mcdowell et al., 2013) thus allowing to record brain data outside of the laboratory over long periods (Hölle et al., 2021). The exciting prospects offered by the exploitation of such mobile research methods have sparked interest in the development of novel signal processing approaches (Reis et al., 2014). Taken together, these developments enable to investigate human cognition directly in naturalistic settings (Ladouce et al., 2017) to tackle fundamental and applied questions across a wide range of research fields such as sport science (Park et al., 2015), architecture (Djebbara et al., 2019) and urban planning (Birenboim et al., 2021), neuroergonomics (Ayaz and Dehais, 2018; Dehais et al., 2020; Gramann et al., 2021), spatial navigation (Thong et al., 2021; Miyakoshi et al., 2021), perception of art and neuroaesthetics (Djebbara et al., 2019; King and Parada, 2021), and the development of assessment and rehabilitation methods for neurocognitive disorders (Kranczioch et al., 2014; Lau-Zhu et al., 2019). As elegantly articulated by Parada (2018), the overarching challenges lying ahead of the MoBI approach to reach its full potential imply a progressive transition from highly-controlled laboratory settings to the study of cognitive phenomena in real-world environments with high ecological validity. Initiating such an incremental approach, a series of influential out-of-lab studies have revisited experimental paradigms commonly used in neuroimaging research and performed them in naturalistic contexts. The following sections present this body of research that established the foundations upon which the present study is based.

### 1.1 Measuring brain activity during motion and outside of the laboratory

In a seminal study, Gramann et al. (2010) revealed that transient brain responses to the presentation of stimuli could be extracted from surface EEG data acquired while participants walked on a treadmill. More specifically, the authors examined the impact of walking speed (standing, walking, and walking briskly) on the P300 component amplitude, which is a widely studied feature of EEG signals whose robustness has established it as a gold standard of EEG research (Polich, 2007). The P300 component is a positive deflection in the time domain of the EEG signal occurring 300ms following the presentation of infrequent or task-related stimuli (typically within the frame of an oddball paradigm) reflective of selective attention processes. These properties established the P300 as a relevant measure to assess the validity and quality of data acquired while subjects were in motion or went outside of the laboratory.

Taking the EEG outside of the laboratory, Debener et al. (2012) demonstrated the feasibility of recording P300 component elicited through an auditory oddball paradigm when participants were walking outside versus sitting inside. The neural responses typically elicited by the presentation of target auditory stimuli were observed in both experimental conditions although they were attenuated in the outdoor-walking condition. In a follow-up study (De Vos et al., 2014), the authors controlled for the environmental factor by having the participants perform the same auditory oddball task while sitting and walking outdoor. Similar to the previous study, an attenuation of the P300 effect was observed for the walking condition which was interpreted as reflecting either a lower signal-to-noise ratio, potentially related to the presence of motion artifacts contaminating the walking data, or a reallocation of attentional resources during walking. These early studies were hinting toward important distinctions between how the mind works under artificial and natural conditions. Circumventing signal-to-noise issues relative to gait-related artifacts, two cycling studies (Zink et al., 2016; Scanlon et al., 2017) confirmed that the attenuation previously reported was partly attributable to the physical activity related to cycling but also, and more importantly, to the higher cognitive demands of being outdoor. Ladouce et al. (2019) further specified the nature of the reallocation of cognitive resources underlying the P300 attenuation by demonstrating that it is not the locomotor demands themselves that take cognitive resources away from the task. Indeed, Ladouce et al. (2019) pinpointed that it is the displacement through space which is substantially taxing in terms of cognitive resources. The reallocation of cognitive resources is therefore due to the increased flow of vestibular and visual information that need to be processed during locomotion. Consistently, Liebherr (2021) reported a reduction of P300 amplitude when students performed the oddball task while finding their way through a university campus as compared to simply walking around a sports field, highlighting the cost of integrating sensorimotor information. Similarly, two studies conducted in real-flight conditions using a passive (Dehais et al., 2019) and active (Dehais et al., 2019) auditory oddball paradigms disclosed lower P300 amplitude when participants faced challenging flying conditions involving an increased flow of visual information to be processed. Taken together, these findings further underlined that cognitive functions are altered when people are immersed in an ever-changing, dynamic and complex environment.

To date, out-of-the-lab studies have mainly resorted to the presentation of artificial stimuli through computerized paradigms (e.g., visual or auditory oddball task), interactions with artificial apparatus taking place within highly controlled environments, or even experimental protocols characterized by the repetition of prototypical behaviours. An important step toward realizing the vision of studying human cognition in the real-world would imply to progressively transition from computerized paradigms to the study of cognitive processes in relation to embodied experiences that are grounded in the real-world. Such a goal remains challenging as the study of cognitive phenomena in the real-world prevents the use of external and artificial experimental probes. In the absence of these experimental markers, questions such as when do cognitive processes happen and how to timestamp and segment the data accordingly become non-trivial issues that need to be addressed in order to accurately extract cognitive events. In contrast to computerized paradigms in which the timing of cognitive events is dictated by the course of the experiment and can therefore be derived from event markers with very high temporal accuracy, the definition of cognitive events in the real-world poses several conceptual and technical questions.

### 1.2 Capturing cognitive events in the real-world

A relevant approach to segment and extract meaningful information from continuous brain recordings is to contextualize the data based on physiological and behavioural data recorded simultaneously. An example of such a multimodal approach can be found in a study carried by Mavros et al. (2016) in which GPS tracking and brain data were combined to study spatial perception and cognitive experiences related to different types of urban environments. Banaei et al. (2017) used Virtual Reality (VR) environments and mobile EEG to study the impact of architectural design on spatial representations when individuals walked through them. Through the use of head orientation information provided by the VR system, the brain signals could be segmented to extract brain dynamics related to the embodied experience of the virtual environment. Another example of contextual information being used to retrieve experimental events from continuous brain imaging recordings is illustrated by Mustile et al. (2021) study in which infrared motion sensors were placed in a real-world environment to detect the passage of individuals over obstacles and extract neural dynamics reflecting motor planning. Hölle et al. (2021) implemented a microphone and a dedicated processing pipeline to detect the onset of natural sounds and lock the associated EEG analyses to unveil auditory attention processes. In a series of studies looking at spatial knowledge acquisition in natural environments (Wunderlich and Gramann, 2021), the experimenter was following the participants navigating through a urban environment and encoded the timing of experimental events manually through a smartphone interface that was synchronized to the EEG system. These manually encoded timestamps, although arguably not of the highest temporal precision, provide nevertheless a basis for the extraction of experimental events. The accuracy in the definition of experimental event timings however has important implications for later EEG analyses. Indeed, the milliseconds scale of ERP components makes their analysis particularly sensitive to temporal lags. The presence of variance across trials in the latency of the responses will likely result in a smearing of the averaged ERP waveform amplitude or in the contamination of ERP components at neighbouring latencies.

In natural settings, first person video recording coupled with eye movements can serve as contextual reference points to segment and analyze the EEG signal to capture neural dynamics of visual processing. Firstly, the video recording can be used to flag and label experimental events in the continuous EEG data by providing information about the onset and end of a visual search task. The eye-tracking data provides a second layer of contextual information to the scene capture, increasing the temporal resolution of the experimental event segmentation. Indeed, the superposition of gaze dynamics to the video recording allow to extract temporal information about the timing of visual events more precisely (Hayhoe and Ballard, 2005). Gaze dynamics are composed of a sequence of discrete fixations whose features (e.g., duration, pattern, previous saccadic distance) provide information about the timing and depth of visual and attentional processes. The timing information of the initial fixations on experimental objects to extract fixation-related potentials (FRP, (Baccino and Manunta, 2005)) has been widely adopted to investigate neural underpinnings of reading and visual search.

Applying FRP analysis approach to free viewing visual search paradigm, Brouwer et al. (2013) demonstrated that the P300 ERP component can be used to infer whether participants are looking at target or non-target stimuli. Kaunitz et al. (2014) also observed the emergence of sensory and attentional components in the FRP associated with the detection of target faces in a scene crowded with distractor faces. The authors further contrasted the free visual search task to a control fixation task and revealed differences in terms of the topography and latency of the P300 component. The free viewing paradigm elicited P300 responses that were most prominent over centro-parietal electrode sites whereas the traditional oddball paradigm exhibited an initial earlier peak at frontal sites followed by later peaks over parieto-occipital sites. These results further demonstrated that the P300 component is robustly elicited upon target detection during visual exploration of natural scenes. Kamienkowski et al. (2012) reported similar FRP components in a free viewing search task and a replay task during which individual elements of the scene are presented as discrete sequences. It is however important to note that in the context of free viewing search task paradigms, participants are typically instructed to keep fixating at the target object when they identify them for an extended period of time. This practice aims to facilitate later ERP analyses by avoiding eye movements artifacts contaminating the EEG signals time-locked to the fixations on targets. Applying such methods, Roberts et al. (2018) and Soto et al. (2018) explored neural markers of faces and economic value processing related to the viewing of visual elements (pictures presented on panels) placed in the real-world using mobile eye-tracking and wireless EEG. In both studies, the panels comprised a central fixation cross to which participants had to return their gaze to after looking at each individual element for a minimum of a few seconds. Despite the artificial nature of the task, these studies demonstrated the feasibility of capturing P300 FRP in a real-world environment. It however remains unclear whether such approach applies in the context of a naturalistic behaviour.

### 1.3 Aim of the present study

The present proof-of-concept study assesses the feasibility of capturing neural markers of visual processing of objects embedded in (i.e., being integral part of) a real-world environment. To achieve this, the present study applies the concept of scalable experimental design (Parada, 2018; Ayaz and Dehais, 2018) according to which similar cognitive phenomena are studied over a spectrum of experiments ranging from highly-controlled to naturalistic environments. Inspired by this approach, the present study contrasts two experimental conditions. The first condition consists in a classic computerized paradigm to elicit neural responses related to the processing of visual information. For this purpose, abstract visual stimuli are presented on a screen and participants are instructed to count the number of occurrence of a certain type of stimuli (i.e., targets). The artificial (as opposed to natural) elements of this condition makes it far removed from experiences taking place in the real-world. In contrast, the second condition maximises ecological validity (i.e., applicability of research findings to real-life contexts) by getting as close as possible to the recording of a natural behaviour taking place in the real-world. To this end, the second condition consists in the performance of a scripted (semi-structured) but realistic behaviour (i.e., searching for a book) grounded in a real-world environment (i.e., a library). Through this leap across the ecological validity continuum, differences related to embodied experiences can be explored while conceptual and methodological gaps pertaining to the study of cognitive processes in the real-world can be uncovered. The present study sets out to address the following questions: Are neural markers of visual cognitive processes elicited through computerized paradigms also present in the real-world? Can such neural markers be captured and their occurrence extracted from the continuous experience of a dynamically ever-changing environment ?

## 2 METHODS

### 2.1 Participants

Twenty-four participants took part in the study. The participants were exempt of any motor, visual and cognitive impairment. All the participants were provided with detailed information regarding the experimental protocol and were introduced to EEG and Eye-Tracking recording procedures. Inclusions and exclusions criteria were checked through the completion of a questionnaire by the participants. The participants gave their written informed consent to take part in the study. The order of the conditions (tablet, library) was counterbalanced across participants to control for potential biases related to fatigue and training. Inconsistencies between synchronization pulses timing sent to the EEG and E-T data streams led to the exclusion of two datasets. The average gaze tracking proportion across participants was above 80 percents for the remaining participants, with most of the missing gaze data coinciding with displacements across the library. Overall these missing data points were therefore not consequential with regards to analyses that concerned time periods during which participants were static (i.e., standing in front of the library shelves). However three additional datasets were excluded from the study due to their insufficient proportion of the Eye-Tracking data containing pupil position (42, 55 and 58%). In those datasets, a substantial portion of gaze data was missing when the participants were scanning through the shelves. In the absence of gaze information at key moments of the experimental paradigm (i.e., when a participant locates a target book cover), it was therefore impossible to retrieve an estimation of experimental events timing. As a consequence of the aforementioned technical issues, only 19 out of the 24 initial recordings were included in the reported analyses.

### 2.2 Paradigm

#### 2.2.1 Tablet condition

Following a standard visual P300 elicitation oddball paradigm, infrequent target stimuli were presented within series of frequent non target stimuli (at a 1:4 ratio). The visual stimuli were presented on a Windows Surface tablet positioned on a library shelf at eye level of the participants. The target stimuli were consisting of red circles while non target stimuli were blue squares of matching areas (circle diameter = 5.1 cm, square length = 4.52 cm). All stimuli were presented in the center of the screen for a 200ms period which was followed by a 800ms interstimulus interval. The participants were instructed to mentally count the number of target stimuli presented. A total of 300 (260 non target, 40 target) stimuli were presented through a Python-based program operating on a 10.8 inches tablet (60 Hz refresh rate, Windows 10 OS). The communication rate of devices and software used to send event markers to the amplifier was measured through blackbox testing (Black Box ToolKit Ltd., York, UK). This approach measures the magnitude and variance of the interval difference between the local timing of stimulus onset on the external device (tablet running the visual presentation program) and the registration of the triggers received on the amplifier end that will be interpreted as timestamps of experimental events in the EEG trace. Accordingly, event marker latencies were corrected to account for the measured delay by subtracting 50 milliseconds (implemented at the beginning of the processing pipeline).

#### 2.2.2 Library condition

The library condition consisted in a natural visual search task and was designed as an analogy of the tablet paradigm previously described. Participants were instructed to search for 40 books in the library. Target visual objects were the books that the participants were instructed to find in the library whereas non-target objects were defined as the four book covers preceding the initial visual exploration of a target book cover 1. Each trial started at a specific location in the library where participants received instructions regarding the location (alley, shelve) of the target book on a sheet of paper that they carried with them.

Several factors were considered for the selection of target books. All the target books selected were placed on shelves that were at eye level of the participants in order to reduce the contamination of the EEG signal by artifacts related to neck movements and other muscular activity. Early works on visual attention have revealed the bottom-up influences of low-level visual features (i.e., intensity, contrast and edge density) on the initial visual exploration of a scene (Peters et al., 2005). The confrontation of computational models inspired by this visual saliency hypothesis with experimental data has later revealed that such bottom-up influences, although partially accounting visual exploration pattern, are however not sufficient to accurately predict visual exploration within the frame of complex scenes (Henderson et al., 2007). Furthermore, empirical evidence from scene perception and visual search experiments highlighted the major contribution of top-down processes (e.g., use of prior semantic information to orient visual search) in how complex scenes are explored and perceived (Underwood, 2009; Birmingham et al., 2009). As both stimulus-driven and cognitive-driven processes interact and influence how information embedded in complex scenes is perceived and processed, several dispositions were taken to ensure consistency in bottom-up and top-down influences across trials. The top-down influence was controlled through the homogeneity of books within a shelve in terms of their semantic field. Indeed, this homogeneity does not favour top-down driven exploration strategies such as parsing and skipping book covers based on prior semantic information gathered (e.g., the target book is more likely to be surrounded by books of the same semantic field). As a consequence of this consideration, shelves with semantically homogeneous books (i.e., related to the same lexical field) were included in the experiment. Moreover, the classification system adopted by the library used as an experimental environment was not based on alphabetical order but followed a systematic catalog arrangement (i.e., books sorted in accordance to the domain and types of publications). This non-alphabetical ordering makes the filtering of content during the initial stage of a bookshelf exploration more difficult. The systematic catalog system could nevertheless be leveraged for a semantic parsing of bookshelf content, but it requires meta-knowledge regarding the classification system used itself and domain-specific knowledge to navigate and parse effectively sections that do not correspond with the title of the target book. The wide variety (genre, types) of the books selected across trials further discouraged the adoption of such domain-specific search strategies by the participants over the course of the experiment. In order to minimize the bottom-up influences of objects whose visual properties make them stand out from the rest of a visual scene, the size and colour of book covers was taken into consideration in the design of the experiment. Indeed, shelves containing books whose covers were particularly salient were not included in the experiment. The position of the book relative to the edge of a bookshelf was another factor taken into consideration for the selection of target books. Indeed, qualitative inspection of preliminary eye-tracking data confirmed that participants mainly scan through the shelves using an initial reading-like approach (left to right and top to bottom) as a default exploration strategy. Therefore in order to gather sufficient fixations on individual books preceding the first fixation on the target book to allow for an analogous analysis, the position of the target book relative to the edges of the bookshelf was purposefully central.

### 2.3 EEG data recording and processing

EEG data was recorded from 32 sensors fitted in an elastic cap following the 10-20 international system which were tethered to a portable amplifier (ANTNeuro, Enschede, Netherlands) recording data at a sampling rate of 500Hz (with a 0.1 to 250Hz online bandpass filter). The amplifier was fitted in an ergonomic backpack carried by the participants. The data was initially referenced to channel Cpz with the ground placed at the Afz electrode site. Electrode impedance was measured prior to each recording session and each channel was maintained below 5 kΩ using electrode gel. EEG data was downsampled to 250Hz and mastoid electrodes (M1 and M2) were discarded.

Continuous EEG data from both recording conditions were processed jointly using the EEGLAB (Delorme and Makeig, 2004) open-source toolbox and custom MATLAB scripts (version R2019b 9.7.0, The MathWorks Inc.). As an initial preprocessing step, the continuous data were visually examined and the portions of the EEG displaying extreme levels of noise (e.g., channel disconnections) were manually discarded. Following this manual data rejection preprocessing, the processing pipeline was divided in two stages. In the first stage, the datasets were filtered with a low-pass filter of 20 Hz and a high-pass filter of 1 Hz with a −6 dB cut-off and a filter order of 1650. Then the continuous EEG was split into consecutive epochs of 1 second. Epochs presenting abnormal values were pruned based on standard statistical criteria (more than three standard deviations from the mean).

Following the initial filtering and removal of noisy data, the first stage of artifact removal was carried out. An extended infomax Independent Component Analysis (ICA, (Bell and Sejnowski, 1995)) was performed on the remaining data, and the resulting Independent Components (ICs) decomposition matrices were saved. In a second stage, the IC features obtained during the first stage of the processing procedure were back-projected to the original filtered data. An automatic classification algorithm (ICLabel) was used to classify ICs (Pion-Tonachini et al., 2019). The results of this classification were examined and ICs identified as artifactual were confirmed manually. The weights of ICs reflecting common artifacts such as eye blinks, eye movements, and heartbeats were subtracted. After this ICA-based data pruning, an average of 58% (SD = 8.2%) of the initial ICs remained across subjects. This proportion of remaining components is in line with the guidelines proposed by (Klug and Gramann, 2020). The ICA-pruned continuous datasets were then epoched around the onset of experimental events (−2000 ms to 2000 ms). Epoched data were then split into the experimental paradigm (tablet and library) and stimulus type (target and non-target) conditions. Averaging across epochs resulted in the obtention of ERP waveforms for each condition. The P300 effect amplitude was computed as the voltage difference (in microVolts) between target and non-target ERPs waveforms within the a-priori time window ranging from 300 to 500ms after stimulus onset. The P300 latency was extracted based on the maximal value recorded within the a-priori time window on a single-trial basis.

### 2.4 Eye-Tracking data recording and processing

Gaze dynamics were recorded using a portable Tobii Pro Glasses 2 eye-tracking system (Tobii Pro AB, Stockholm, Sweden). E-T data was acquired from four eye cameras tracking pupil position and corneal reflection binocularly at a sampling rate of 100Hz. Built-in parallax and slippage compensation methods were performed to maintain E-T tracking accuracy during movement. The calibration procedure consisted in presenting a target placed at 1, 3 and 5 meters distance from the participants to ensure a reliable tracking at different fixation depths. The eye-tracking apparatus comprises the camera-equipped glasses and a recording unit to which the glasses were connected through HDMI. The recording unit was fitted in the backpack with the mobile EEG amplifier to which it was connected through micro USB to a 8-bit trigger input. A timestamp was generated by the E-T every 5 seconds and sent to the EEG amplifier for synchronization purposes. Prior to the study, the accuracy of the synchronization triggers has been extensively tested over hours long recordings. The delay between recording systems remained below 10ms and was consistent throughout the testing recordings. The raw E-T data was then reviewed visually and time periods characterized by poor tracking accuracy were recalibrated using known fixation points (e.g., participants were instructed to look at a fixation point at the beginning of each trial). Missing gaze samples were interpolated (using a moving median of 5 samples) if the gap between retrieved samples was lesser than 75ms, otherwise the samples were considered lost. The proportion of gaze samples retrieved throughout the recording (expressed in percentage) was over 75% for the majority of participants. As mentioned above, two outlying datasets had to be excluded due to their low proportion of E-T samples recorded. The continuous data was then subjected to a noise reduction function based on a non-weighted moving median filter with a window size of 3 samples. A classification algorithm was then performed on the raw E-T data to identify fixations. The built-in Tobii I-VT Fixation Filter was used with a velocity (expressed in visual degrees per second) threshold of 30°/s over 20ms window length. Gaze samples above the velocity threshold were classified as saccade samples. Short fixations lasting less than 50ms were discarded. Adjacent short fixations were merged when their inter-fixation (saccade) duration was lower than 75ms or that the visual angle difference between these fixations was lower than 0.5°. Henderson and Luke (2014) have reported that the mean fixation time was around 250ms during complex scene visual search tasks and that this fixation duration, although prone to inter-subject variability, was stable within and between sessions. Based on these findings a lower threshold of 200ms was used for the definition of visual fixations.

### 2.5 Event extraction

The processed E-T data was then visually inspected by the experimenter and experimental events timing were manually annotated using the Tobii Pro Lab software. The onset of the initial fixation on a target book cover was used as a timestamp for the definition of a target trial. The timing of the onset of preceding fixations on four distinct book covers were used to retrieve non-target experimental events. This approach to the definition of the library experimental events timing was adopted to allow for comparisons with the tablet condition in which target stimuli were presented in the midst of non target stimuli with a 1:4 ratio.

### 2.6 Statistical analysis

Statistical analyses were performed on the mean amplitudes within the P300 time window (300 to 500ms) recorded at Pz electrode site where the P300 is most prominent (Alexander et al., 1996; Polich et al., 1997). Repeated measures ANOVA and paired-samples t-tests were performed on the extracted amplitude features. To ensure that parametric analysis was appropriate, a normality test was carried to ensure that the data followed a normal Gaussian distribution. In addition, Holm-Bonferroni correction for multiple comparisons was applied for all post-hoc t-tests.

## 3 RESULTS

### 3.1 ERP analyses

A repeated-measures ANOVA was carried on ERP features with the experimental paradigm (tablet, library) and stimulus type (target, non-target) as factors. Post-hoc paired-sample t-tests were carried to explore main effects.

#### 3.1.1 P300 ERP

The repeated measures ANOVA revealed that both the paradigm [F(1,18) = 8.98, p < 0.01, n2p = .33] and the stimulus type [F(1,18) = 17.62, p < 0.001, n2p = .49] had a main effect on P300 ERP amplitude. Moreover an interaction [F(1,18) = 19.98, p < 0.001, n2p = .52] between the two factors was found.

Post-hoc comparisons revealed that target stimuli elicited P300 ERP responses of significantly higher amplitude than non-target stimuli for the tablet condition [t(18) = 5.2, p < .001, d = 1.19, BF10 = 450.9] but not for the library condition [t(18) = 1.33, p = .209, d = .29, BF10 = .49]. The present results indicate that a P300 ERP response was consistently elicited in the tablet condition whereas the effect was not observed in the library data (as illustrated in Figure 2).

**FIGURE 1.**
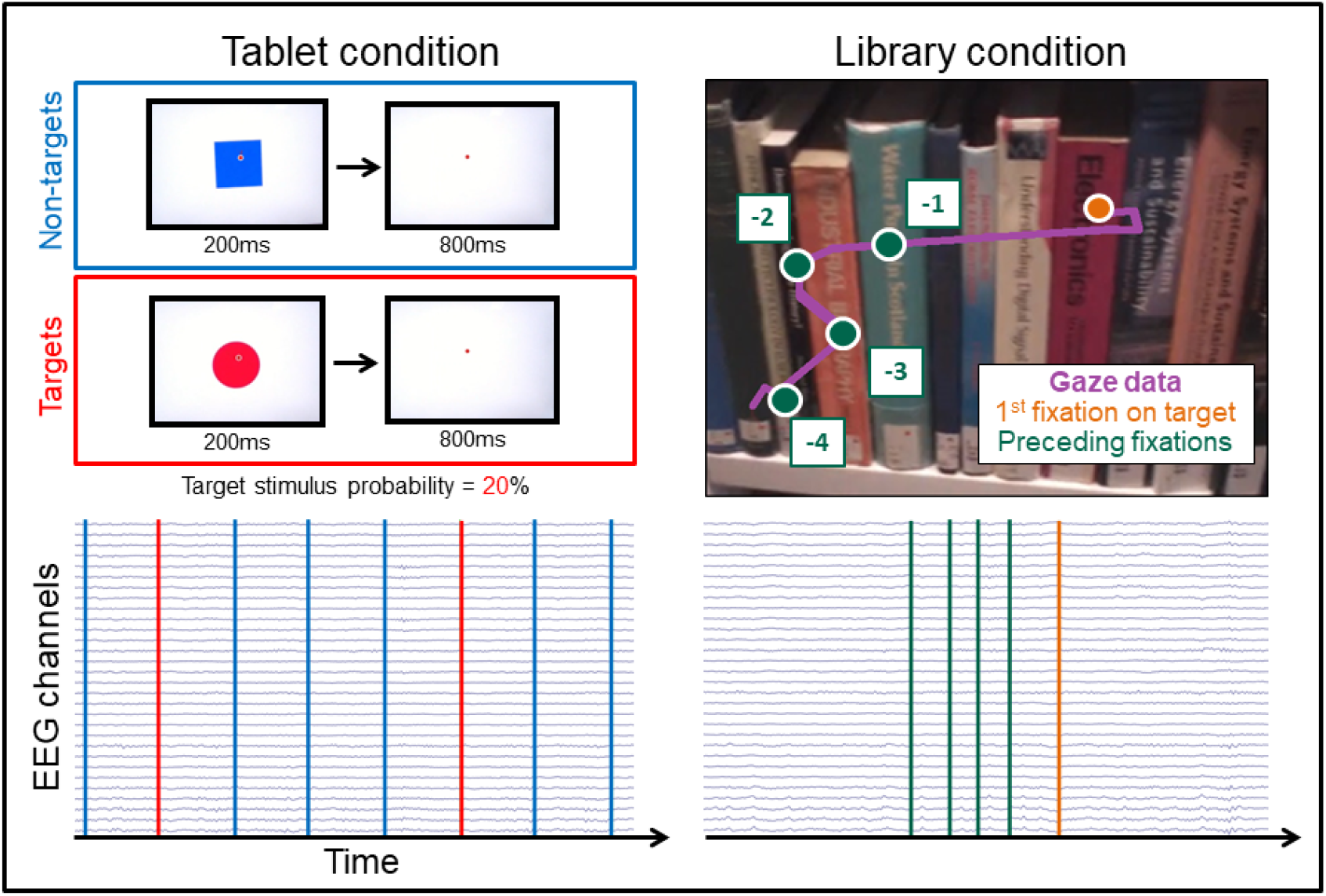
Example of the two types of experimental events (target and non-target stimuli) for both conditions (tablet on the left and library on the right side). For the tablet condition, series of discrete visual stimuli were presented through a computerized paradigm. The infrequent target stimuli were red circles and the frequent stimuli were blue squares (1:4 ratio). The stimuli were presented for 200ms followed by a 800ms interstimulus period. The onset of visual stimuli was used to define experimental events timing. As can be observed from the continuous 32 channels EEG data, event markers were equally spaced over the time series. For the library condition, gaze data and first person video recording were reviewed by the experimenter. The experimental event timing was defined as the onset of a fixation on the target book cover for target trials and as the first fixation on the four preceding book covers for the non-target trials. The variability in terms of fixation and saccade duration results in an irregular temporal distribution of visual events as denoted by the lower right plot.

**FIGURE 2.**
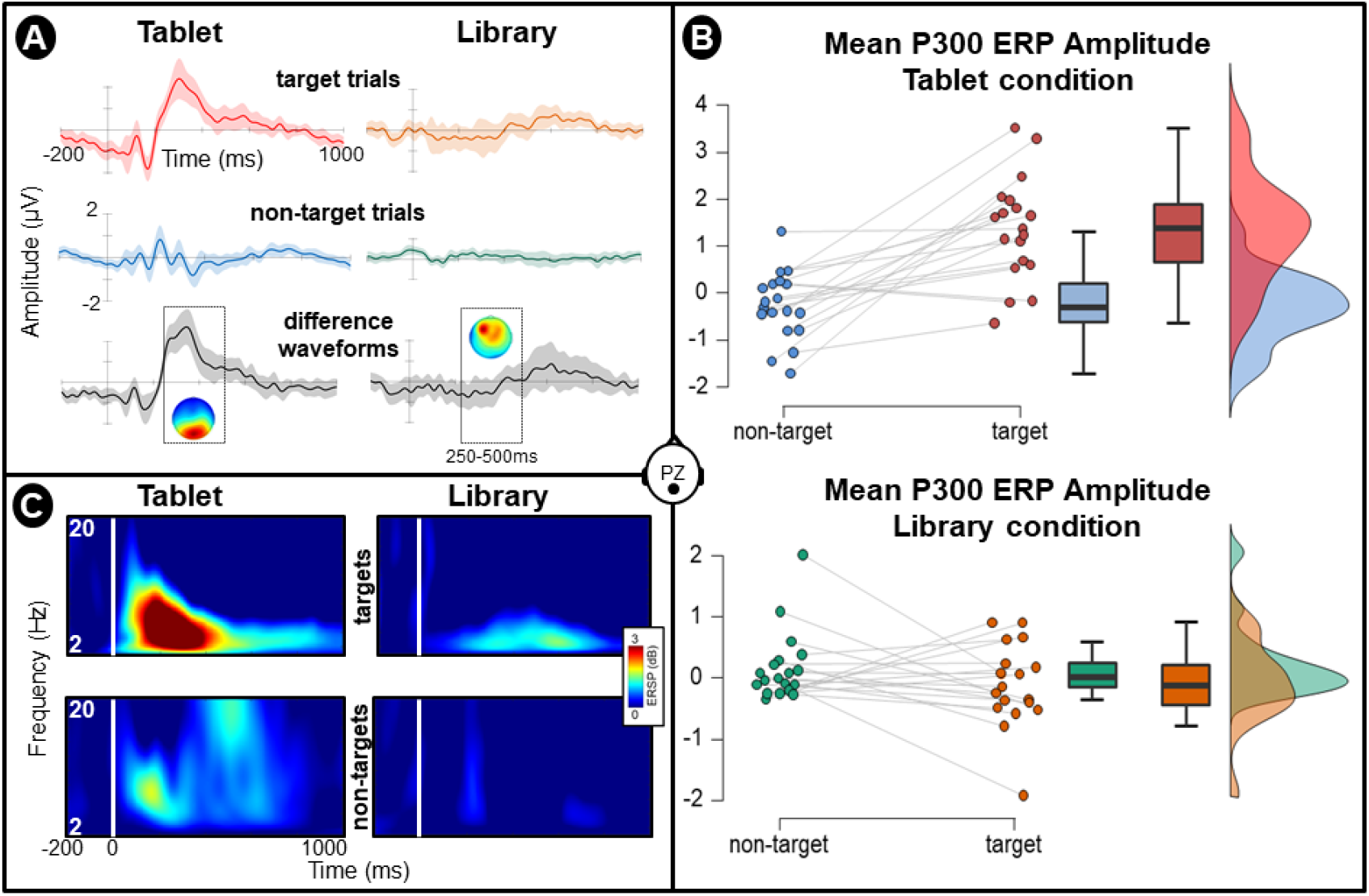
Grand average (N=19) Event-Related Potentials recorded at parietal electrode (Pz). A. Grand-average ERP waveforms elicited by target and non-target visual stimuli for both tablet (left) and library (right) conditions. The difference ERP waveform is plotted below with a scalp map presenting the topographical distribution of activity recorded within the P300 time window (250 to 500ms). B. Distribution of mean P300 ERP amplitude recorded at Pz electrode across participants. C. Time-frequency plots presenting mean event-related changes in spectral power (deciBel scale) elicited by target and non-target stimuli for each experimental condition (tablet and library).

### 3.2 Interim discussion

As discussed in the previous sections, there are many unknown variables affecting the definition of the onset of a cognitive event in the real-world. The a-priori approaches based on gaze data (i.e., fixations on experimental objects) may not coincide with the actual onset of cognitive processing of that particular visual information. The cognitive processes related to a visual fixation may precede or follow the initial fixation. The temporal gaps in either of those scenarios would introduce variance in the latency of ERP responses which would not survive averaging processes. The inspection of gaze data provided striking evidence that both scenarios (i.e., fERP based definition of visual processes onset being late or early) were commonly found within single recordings and across participants.

Indeed, the first fixation on the target book may already be relatively late with regards to the overall temporal course of the visual processing of that information, which may have already started when the information entered the peripheral visual field. This possibility is illustrated by the gaze pattern of the participants preceding target identification: the visual exploration strategy typically shifts from an orderly scan of the book covers to a sudden ballistic saccade toward the target. The large angular distance of these saccades further suggest that the book covers present in the periphery are already being processed semantically. In this scenario, the ERP responses related to the visual processing of these target trials would precede the initial fixations and therefore the ERP would be temporally shifted in time in what is commonly called the prestimulus period. This shift is particularly problematic for the application of baseline correction approaches that aim to detrend the data using this prestimulus period. If the prestimulus period contains the signal of interest, then subtracting it from later time series signals would introduce antagonist artifactual effects.

In contrast, all recordings contained several trials in which the individuals visually explored the target book cover without identifying them as targets and continued their exploration of the shelves. The object may have only been identified as a target after having been fixated on. In that second scenario, using the fixation on the object as a timestamp for the onset of visual processing introduces a delay in the responses shifting the neural signals at a later point in the time series. This second scenario is more likely to occur when the target object is less salient and/or that there is more competing visual information in the visual field such that the bottom-up influences of the non-target objects counteract top-down strategies. Although these “missed” trials were relatively infrequent (their number was too low to allow for a dedicated ERP analysis), averaging them with actual target identification trials would lower the SNR of ERP responses of the latter. The strict definition of first fixation on a target book cover was therefore relaxed and the fERP onset for such trials was changed to the first fixation on the target preceding their actual identification (i.e., participant stops scanning the shelves and return to the starting point).

Both of these phenomena are likely to occur in real-world environments where the objects and their surroundings visual features are variables. Not only the cover of the target books may have been more or less salient, but the density of the books on the shelves, visual properties of the books’ covers (i.e., colours, width, orientation), the lighting in different places of the library and other priming effects through semantic association (e.g., shared lexical field) induced by surrounding cover titles are as many factors that may affect the timing between early visual and later attentional processing of target objects and their visual exploration. Therefore, the naive fixation-based approach for the timestamping of experimental events is inherently limited to capture accurately the onset of cognitive processing of visual information embedded in the real-world.

Another source of temporal imprecision comes from the relative low sampling rate of the Eye-tracker scene camera. Indeed, the scene recording is captured at a rate of 30 frames per second. There is therefore a 33ms gap between every frame. This gap means that the accuracy of visual events timing achieved by reviewing the video recording frame by frame is limited by the temporal resolution of the scene capture. In addition to this temporal variance, the gaze data superposed to the scene recording is acquired at a higher rate (100Hz) and is smoothed in order to match both recordings. Considering the additional degrees of freedom that apply to a fully mobile eye-tracking system, it is sensible to assume that the matching between the scene recording and gaze data points may be subject to some imprecision, especially during head movements and even more so during whole body movements. Any incoherence between the data streams comes however at the price of an additional 33ms variance added on top of the original 33ms. The milliseconds scale of ERP components and the averaging process usually applied to uncover the signal from background activity make them particularly sensitive to subtle temporal variations. Taken together, the aforementioned considerations suggest that mobile eye-tracking data, while providing contextual information for the definition of experimental events, may however not provide a temporal estimate sufficiently precise to perform ERP analyses. The direct consequence of any of these source of temporal imprecision is that there would be important variations across trials in terms of ERP latencies, essentially leading up to smearing effects (Ouyang et al., 2011) or even cancelling out potential ERP components through averaging procedures.

While it appears plausible that the absence of ERP components time-locked to the initial fixation on experimental objects observed in the library condition may be caused by inter-trial latency variability, it is nevertheless important to consider that such brain signals may simply not be present during the library visual search task. Indeed, the P300 ERP component could be an artificial response evoked by computerized paradigms that do not transfer to the realworld. The former hypothesis implies that ERP components would manifest at single-trial level but would be shifted in the time-domain whereas the latter hypothesis implies the total absence of such ERP components for the library condition. In order to elucidate these competing assumptions, single-trial ERP responses to target book covers were inspected over a larger time around their fixation onset. As can be observed in Figure 3, signals sharing spatial (parietal topographical distribution) and spectral features (delta and theta band activation) of the P300 ERP are present at single-trial level. The variance in the latency of such signals is however important, spanning across both the prefixation and post-fixation periods, with a wide temporal distribution over the latter. These observations suggest that the P300 ERP response may be present in the library condition recordings but have substantial variance in their latency. The fixation-related potential approach applied to define the onset of cognitive event may not be valid in the context of the present real-world data. It remains unknown whether the P300-like signals observed at single-trial level is effectively a specific response to target stimuli. To address this question, library epoched data of both target and non-target stimuli were compared to a subject-specific template of P300 ERP response based on the tablet ERP average waveform. A higher similarity between target waveforms than between the template and non-target stimuli would provide further evidence suggesting that the library target responses are effectively reflecting time-shifted P300 ERP responses.

**FIGURE 3.**
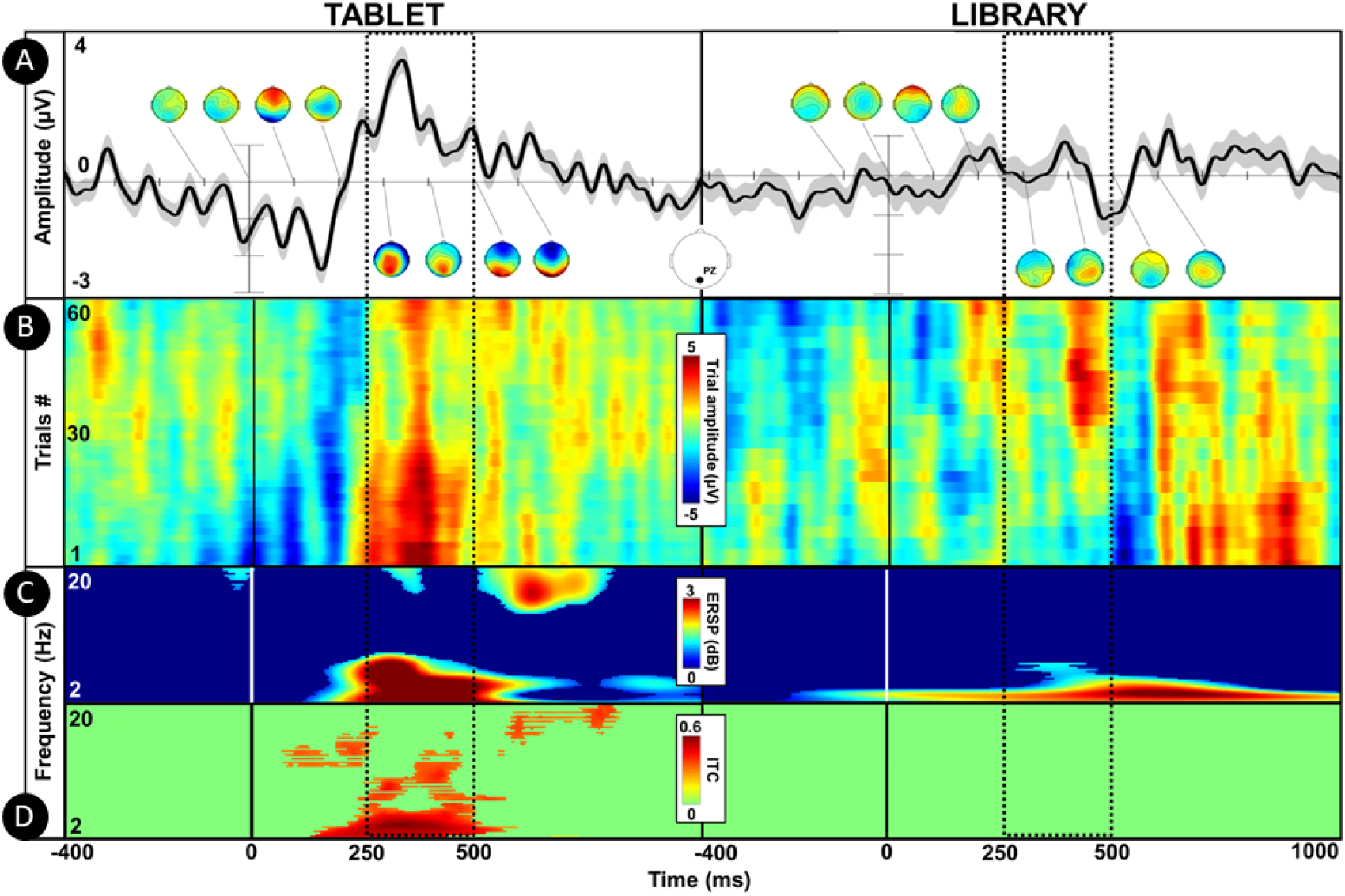
Illustrative single subject (Participant 5) fERP and ERP channel data (Pz electrode) comparing temporal dynamics across the library and tablet conditions. The dotted windows indicate the a-priori time window commonly used to extract P300 ERP features in experiments using computerized paradigms (250 to 500ms). A. The uppermost plots present the grand average ERP waveforms around the onset of experimental events which were defined as the first fixation on the target book cover for the library condition and the onset of stimulus presentation for the tablet condition. A series of topographical scalp maps indicate the spatial distribution of ERP responses over time (from −100 to 600ms around event onset, 100ms incremental step). B. ERP-image plot presenting single-trial broadband ERP responses stacked over the y-axis. This plot reveals a delayed amplitude increase in the case of the tablet condition with respect to the prototypical P300 ERP response latency observed that can be observed in the tablet condition. C. Average time-frequency activity relative to baseline period (−200 to 0ms) in deciBels (dB). The P300 ERP response is mainly arising from delta and theta-band (1-8Hz) activity. D. Inter-Trial Coherence (ITC) measure of time-frequency dynamics. A statistical mask with a alpha of 0.05 was applied (permutation t-tests, FDR correction). The tablet condition exhibits a strong coherence across trials within the P300 ERP time window and frequency range whereas no coherence is found in time-frequency dynamics for the library condition. Taken together these results highlight that the tablet condition reliably evoked P300 ERP responses that were time and phase-locked to the onset of stimulus presentation. In contrast, the library condition is characterized by EEG signals that share P300 characteristic features but that are distributed over a wide time range.

### 3.3 Assessing similarity between library and tablet signals

A template matching method accounting for temporal shifts is required in order to assess whether the P300-like signals observed across the library EEG data are similar to time-locked ERP responses recorded in the tablet condition. Dynamic Time-Warping (DTW) algorithms allow to compute measures of similarity between time series that may vary in speed and consequently be shifted in time. DTW is an ubiquitous approach commonly applied to speech recognition to handle variations in speaking speed. DTW algorithms compute an optimal match between two time series, in the present case a template based on the average ERP waveform of the tablet condition and single-trial ERP waveforms of the library condition. The sequences are non-linearly warped (i.e., shifted) in the time domain, every index of each sequence being matched with at least one index from the other time series. A cost measure is computed as the sum of absolute distance values of each matched pair of indices. The optimal match is selected on a minimal cost basis. The similarity (referred as dissimilarity depending on applications) measure provided by DTW accounts for amplitude differences between the signals following the non-linear warping.

For each participant a template was defined as the average ERP waveform (1 to 8Hz bandpass filtered) of target stimuli during the tablet condition (see Figure 4 A). DTW similarity measures were computed between the template and single-trial time series (1 to 8Hz bandpass filter) for both target and non-target stimuli of the library EEG data. The library target trials were significantly more similar to the template than the non-target trials [t(18) = 4.58, p < .001, d = 1.05, BF10 = 132.64] following non-linear warping in the time domain. This similarity further support that P300 ERP responses are present in EEG data epoched around the first fixation on the target book covers as these time series are more to typical P300 ERP waveform elicited by a computerized paradigm.

**FIGURE 4.**
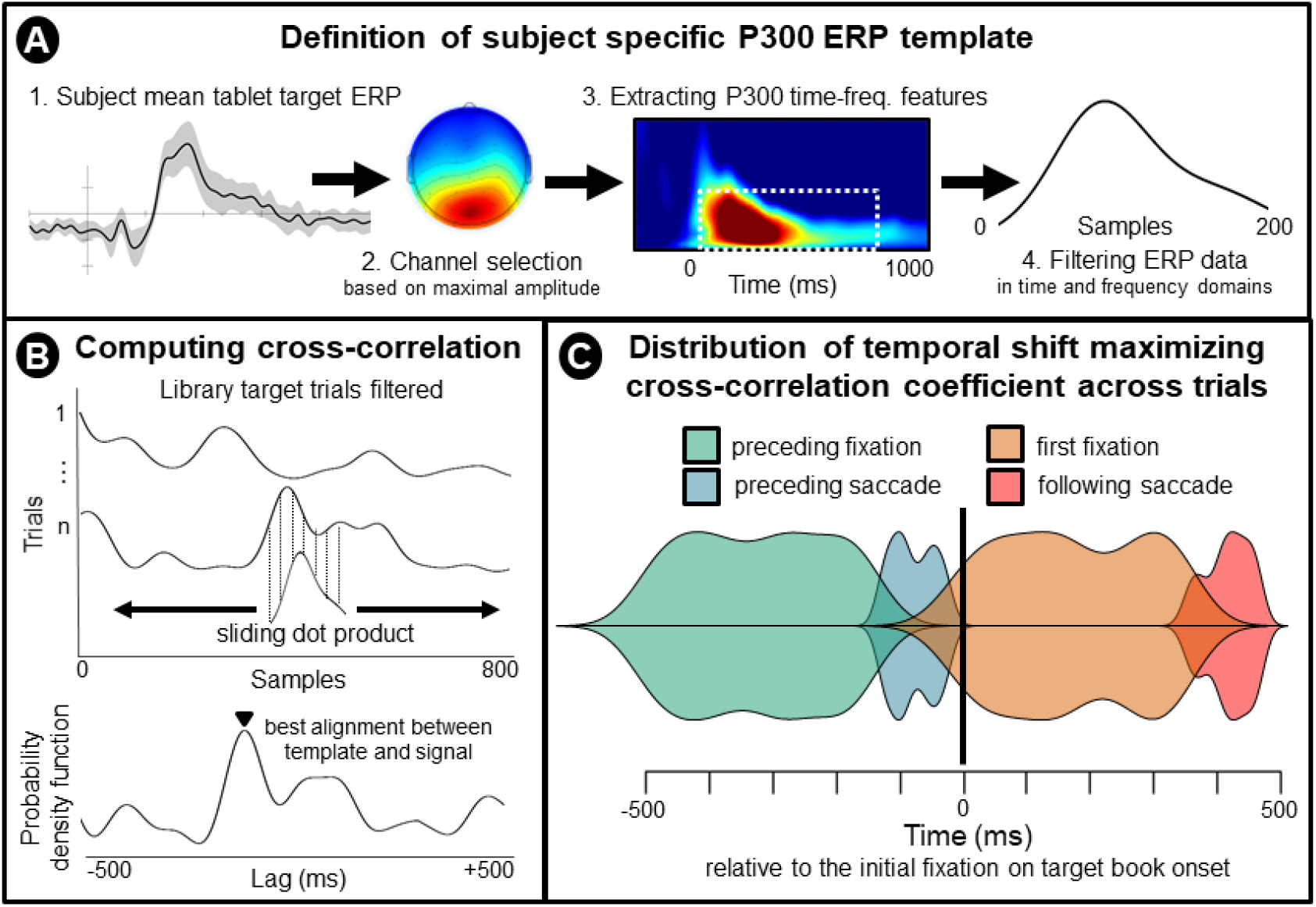
Implementation of cross-correlation measures for the time-domain alignment of P300 ERP responses recorded in the library condition. A. First, a subject specific template was computed as the mean ERP response elicited by target trials of the tablet condition. Secondly, the channel exhibiting the maximal ERP amplitude within the P300 time window (250-500ms) was selected. Third, a time-frequency decomposition of the ERP was then performed on the selected channel data, the frequencies and temporal features of the P300 ERP responses were extracted. Lastly, the selected channel data was then filtered at the frequencies contributing the most to the P300 ERP and data points included in the temporal window previously defined were then used to create a filtered P300 ERP template. B. The library target data was filtered accordingly to the spectral and spatial filters applied to the template data. A cross-correlation between the template and the signal data was performed for each target trial of the library condition. The resulting sliding dot product was used to create a probability density function whose maximal value indicate the best alignment between the template and the signal. The lag associated with the maximal value was then used to correct the single trial onset timing. C. Distribution of the time correction applied across all library target single-trials and the corresponding gaze dynamics occurring at the time (relative to the first fixation on target books).

### 3.4 Alignment of real-world ERP responses

The previous observations and analyses have provided evidence that P300 ERP features are present around the first fixation on target book covers but are not time-locked to the fixation timing. In order to perform ERP analyses on the library data it is critical to address the latency variability of its ERP responses. The cross-correlation method computes the similarity between two time series that are shifted along each other. The result of this convolution is a sliding dot product whose maximum value informs about the lag between the two series that optimizes similarity (see Figure 4 B). This method is useful to search for known features within long signals. Following a similar implementation than DTW, a subject-specific template of the tablet P300 ERP is slid over every library EEG data epochs. The temporal lag maximising the similarity between the time series will then be used to align the library single-trial data. The average temporal lag between the tablet P300 template and library single-trial data was −80ms (SD = 344ms). The first fixation on target stimuli timing used for epoch extraction was corrected on a single-trial basis. The continuous EEG data was then epoched around lag corrected markers. The following sections present ERP analyses performed on time-corrected epoched data.

### 3.5 Lag corrected ERP analyses

#### 3.5.1 P300 ERP Amplitude

The repeated measures ANOVA revealed that the condition did not have a main effect on P300 ERP amplitude [F(1,18) = 0.44, p = 0.51, n2p = .007] anymore while the stimulus type [F(1,18) = 16.82, p < 0.001, n2p = .27] had a main effect on P300 ERP amplitude. There was no interaction [F(1,18) = .996, p = .33, n2p = .007] between the two factors on P300 ERP amplitude. Post-hoc comparisons revealed that target stimuli elicited P300 ERP responses of significantly higher amplitude than non-target stimuli in the lag corrected library condition [t(18) = 2.403, p = .027, d = .551, BF10 = 2.289]. Interestingly, no significant difference was found in P300 ERP amplitude between the lag corrected library and tablet conditions [t(18) = .454, p = .655, d = .104, BF10 = .261] as can be observed in Figure 5.

**FIGURE 5.**
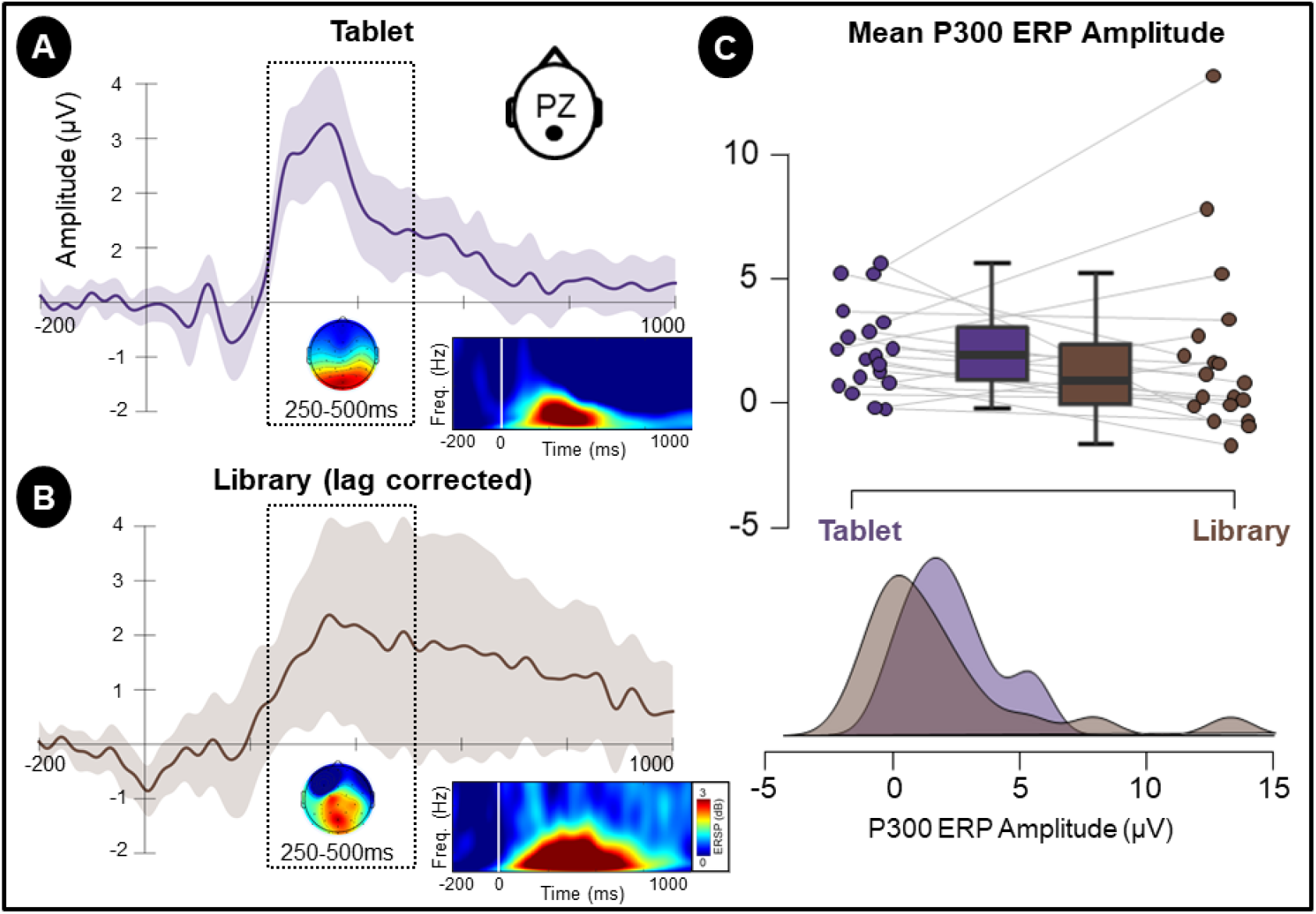
Grand average (N=19) Event-Related Potentials difference (targets - non-targets) waveforms recorded at parietal electrode (Pz). A. Grand-average ERP waveforms for the tablet condition. The topographical distribution within the P300 ERP time window (250 to 500ms, see dotted frame) is presented as a scalp map. A time-frequency plot presents mean event-related changes in spectral power (deciBel relative to −200 to 0ms prestimulus baseline). B. Grand-average ERP waveforms, scalp map and time-frequency plot for the library condition after lag correction. C. Graphical visualizations of Mean P300 ERP amplitude distribution across experimental conditions.

#### 3.5.2 P300 ERP Latency

There was no significant difference found in P300 ERP peak latency between the tablet and library conditions [t(18) = 1.371, p = .187, d = .314, BF10 = .532] after lag correction was applied to the library data.

## 4 DISCUSSION

The present study was designed to assess the feasibility of capturing neural markers of visual attention related to the processing of objects embedded in the real-world. Inspired by the scalable experimental design approach, two conditions at both ends of the experimental control and ecological validity continuum were contrasted: a classic visual oddball paradigm running on a tablet and a naturalistic visual search task of book covers in a library. While the tablet paradigm presented series of discrete stimuli whose onset was timestamped in the EEG data, the library EEG data was epoched around visual fixations on experimental objects. In the initial analyses the presence of the P300 ERP response was only found for the tablet condition. The inspection of single-trial data recorded in the library revealed signals whose features were similar to the P300 ERP found for the tablet condition but those signals were shifted in time. However, this raises the question on how to determine the timing of the processing of the visual objects (i.e books) embedded in the real-world. Such a definition is highly dependent on the validity of the measure that has been used. Should that definition be based on the visual information entering the field of view of the individual already raises issues regarding the definition of this visual field. Is the phenomenological experience of a visual object bound to its entrance into the foveal spotlight? In this case then the initial fixation on a visual object appears a valid approach to extract brain dynamics reflective of a visual cognitive experience. There is empirical evidence from reading research on text processing that visual information present in parafoveal areas are already processed at sensory but also at semantic levels (Pan et al., 2021). Using the entrance within the parafoveal field of view to define the onset of visual event requires computing the angular distance between the individual’s retina and the visual object. In the absence of a specific continuous measure of such distance over the course of the experiment, we performed DTW similarity measures using individual subject template of the P300 response that was based on the tablet data and signals recorded in the library. The similarity measures with the tablet template was substantially higher for signals around visual fixations on target book covers than non-target book covers, suggesting the presence of time-shifted P300 ERP that are specific to target trials in the library condition. While DTW is a powerful method to evaluate the likeness of ERP responses between library and tablet conditions, it does not provide a measurement of how much time series have to be warped in the time domain to match the template. Moreover the non-linear warping may introduce distortions of the time series and the resulting warped series may be of different lengths (i.e., different number of data points) which may bias further statistical analyses (Zhao et al., 2020). Another approach was therefore needed to quantitatively characterize ERP features while accounting for temporal lags. The cross-correlation coefficient was therefore used to find the lag that would maximize the similarity between the tablet template and the library data upon which target events latency were then lag corrected. The realignment of the library signals uncovered a P300 ERP response. The present findings suggest that neural markers of visual attention elicited by computerized paradigms are also present in a naturalistic environment. These results demonstrate the potential of the joint recording of EEG and Eye-Tracking data to study visual cognition in the real-world. Moreover, retrieval of cognitive events latency through analyses carried upon EEG data can in turn be used to gain insight about gaze dynamics involved in object recognition and processing (see Figure 4 C). These results also highlight the relevance of data-driven approaches such as template-matching to recover neural signals of interest in real-world data. However these findings should be interpreted cautiously. For instance, did the P300 evoked response account for the same cognitive phenomena in the lab and real world scenarios? Is it possible to compare the amplitude and the latency of this latter ERP across the two experimental conditions and draw conclusion about the effect of environmental factors and task difficulty previously reported in several studies (Cortney Bradford et al., 2019; Ladouce et al., 2019; Dehais et al., 2019). The following sections will address critical differences between the two experimental settings (tablet and library conditions) and discuss the implications of such discrepancies.

### 4.1 Differences between free-viewing and gaze-fixed paradigms

In a gaze-fixed scenario such as the tablet paradigm, no particular predictions of information localization nor eye movements are required. In contrast, the active free-viewing search task in the library implies an exploration of the visual environment in order to identify the identity of a stimulus (target or non target). The decision of where to look next at any given time is therefore critical. Due to the high degree of freedom inherent to real-world settings such as in the library condition, our participants’ cognitive experiences were more complex, dynamic and multidimensional. Indeed, at any given time, there is a wide range of multimodal sensory information to process, with each of these information having a certain number of potential states. Based on this information input, the nervous system works towards building an understanding of the present state of the environment and attempts to predict future states. These differences have several implications. First, that the higher order cognitive processes involved in the free-viewing library task to update contextual information and guide the visual exploration of the environment do not correspond to those involved in the gaze-fixed tablet condition. Secondly, the EEG data recorded in a free-viewing task inherently includes both neural activity related to experimental events but also ocular artifacts (especially within the frame of FRP analyses, see Ehinger and Dimigen (2019)). The temporal overlap between neural responses evoked by consecutive fixations is another critical issue related to free viewing conditions. Indeed, the series of rapid eye movements that typically precedes the initial fixation on an experimental object may all bring their respective neural signatures which could be confounded for the visual event-related potentials as their delayed latencies and their short temporal spacing may overlap with the latter. While later components such as the P300 are less prone to this issue than early sensory processing components (i.e. N1 and P1 Dandekar et al. (2012)), it is worth noting that regression-based analyses approaches have been proposed to disentangle the contribution of consecutive eye movements on EEG signals based on eye-tracking information (Dimigen and Ehinger, 2021). Furthermore, the presence of neighbouring visual information requires increasing cognitive demands to select the target information and inhibit the exploration and processing of concurrent information (Hillyard et al., 1973). In the case of the free-viewing library task, top-down cognitive processes direct attentional resources towards the detection and identification of target while inhibiting bottom-up influences of distracting information. The nature of this competition for attentional resources does not apply to the same extent to the tablet condition.

### 4.2 Permanency and uniqueness of real-world visual experiences

The perceptual differences between target stimuli in the experimental tasks is another factor that could lead to differences in terms of neural responses recorded between the tablet and library conditions. It has been largely documented in traditional P300 experiments that a higher contrast between target and non target stimuli lead to an increase of P300 amplitude. The saliency of target stimuli has been shown to largely contribute to this effect (Luo and Ding, 2020). The tablet experiment uses variations of low-level perceptual properties of the stimuli such as shapes and colours to make the target and non target highly distinguishable from each other. The target book covers used in this experiment were all different, and so were their respective neighbouring book covers. More than shapes and colours, the shelves density, and the width of covers systematically varied across trials recorded in the naturalistic library setting. This means that each trial was singularly different from another in terms of how target and non target stimuli were contrasted and how relatively salient the target stimuli were. The objective of the present study was to dive directly into the investigation of neural dynamics during free visual exploration of a real-world environment. For this reason, the experimental setup took place in a naturalistic environment that was not altered by the introduction of experimental probes. Nevertheless, the additional degrees of freedom pertaining to the use of various books as target stimuli in the library condition could however be circumvented through the use of identical book covers that would be placed in the vicinity of other book covers that would share similar low-level perceptual properties. While such an approach is not an ideal solution to achieve the vision of studying cognitive experiences directly in the real-world and may be impractical to setup, it could however help reconcile the measures by reducing potential differences related to lowlevel visual processes between experimental conditions. Eventually the permanence of the visual information is yet another difference between the two experimental conditions. What is meant by permanence here is that the visual objects are always present in the environment, before and after being experienced by the user. This is in contrast to a discrete visual event such as a sudden flash of light or images appearing on a monitor. In the latter cases, the phenomenal visual experience can be timely defined in coincidence of the onset and offset of the physical apparition of the visual stimuli. A distinction should nevertheless be made between experimental event and cognitive phenomenon. Computerized paradigms use the former to elicit the latter. In the real-world, how such cognitive phenomena are expressed and how they can be captured remains unknown. Therefore, a sensible question is whether neural responses typically observed in the laboratory such as the P300 ERP are merely just a by-product of the specific way sequences of stimuli are presented and are therefore not present in everyday life experiences.

### 4.3 Considerations regarding brain and body imaging in the real-world

#### 4.3.1 Template Matching approaches: advantages and limitations

This study clearly illustrates how capturing neural responses to visual objects embedded in the real-world constitutes a substantial leap (both technically and conceptually) from recording evoked potentials elicited under highly controlled laboratory settings. These challenges were addressed through a combination of novel methodologies ranging from the experimental design to the recording and analysis of the data. However, the template-matching approaches (DTW and cross-correlation) applied to correct the library data event latencies presents several pitfalls that are crucial to consider in order to interpret the results. First, by comparing the library EEG signal recorded few seconds before and after the first fixation on the target with a prototypical signal representing a P300 effect recorded using a computerized paradigm, a strong assumption regarding the expression of the visual cognitive event in the library is made. While this assumption is directly derived from the original hypotheses of the study, and therefore provide a sound basis for the application of such method, the results should nevertheless be interpreted with caution. Indeed, by identifying features in the library data that resemble the template, the alignment resulting from the correction may produce a waveform whose features are similar to the template but whose origin may be different (i.e., brain signals not related to the task or even noise). While this possibility cannot be entirely dismissed, it should be noted that the relative proximity of the corrected responses (indicated by the low lag values around the original fixation point), and the presence of other components that were not included in the template and its localization (as indicated by topographical maps) together suggest that the latency-corrected signals share properties with the tablet signals that go beyond mere spectral and temporal domains similarities. Second, the application of such approach to the library data not only provide a mean to correct the latency of the effect (provided that the previous assumption is valid) but this approach also artificially reduces the temporal variance of such signals in the corrected data by imposing strong temporal constraints which are based on the template data. This issue has implications for the interpretation of both the amplitude and latency of the P300 ERP. Indeed, the strong time-locking and phase locking induced artificially by the realignment of the EEG signals will exhibit minimal temporal variance which may inflate the amplitude of the averaged waveform as it is not subject to the same smearing that would naturally occur in relation to variance in the P300 latency across trials. Moreover, variations in P300 latency may be particularly informative within the frame of real-world experiments as several aspects of the visual stimuli (i.e., book cover more or less salient) but also environmental factors (i.e., competition of surrounding visual information) could have an impact on cognitive processing speed. Although the cross-correlation based template matching approach is not an ideal solution, the corrected latencies derived from the EEG signal template provide information that open a new range of analyses to be carried on the multimodal data.

#### 4.3.2 Intrusiveness of MoBI devices: toward transparent solutions

The participants were equipped with a research-grade mobile EEG system that comprised an EEG cap tethered to an amplifier which was fitted in a backpack and eye-tracking glasses also tethered to a recording unit. While the eye-tracking glasses have been deemed as relatively unobtrusive following a short adaptation period, wearing the EEG cap in a public space such as a university library was reportedly a self-conscious experience for the participants. While data quality is a critical for the selection of EEG systems, the degree of comfort and discretion are also key aspects to consider in the context of real-world studies. An elegant solution to this issue could be found in minimalist EEG devices. For example, around-the-ear electrode arrays are discreet, quick to setup and can be worn comfortably for extended periods of time (Debener et al., 2015). The minimalist and ear-centered montage does however not offer the spatial resolution of a whole-head EEG system and therefore may not be sensitive to far-field potentials (Meiser et al., 2020). Nonetheless around-the ear arrays have proven to be an effective solution to record neural markers of visual processes (Pacharra et al., 2017) and selective attention to auditory information (Debener et al., 2015) (Mirkovic et al., 2016) (Bleichner et al., 2015). This comfortable solution is particularly appealing for the study of everyday-life human cognition but also offers promising medical use cases such as sleep staging and long-term monitoring of epileptiform activity (Bleichner and Debener, 2017). The future of MoBI research field is linked to the development of inconspicuous and comfortable recording devices as they will enable the range of research to expand to social interactions in real-life situations.

#### 4.3.3 Ethics of real-world brain and body imaging data

A wide range of personal information is gathered from pervasive devices (i.e., mobile phones, smartwatches). The recording of GPS, heart rate, and accelerometers data can be used to track where and what an individual is doing at various times of the day. Although privacy concerns are raised sporadically, the quantity and variety of data collected from individuals has been steadily on the rise. More pointedly, instrumentation that was limited to medical and research purposes are now ported into consumer-grade devices. As this new generation of devices find their way to households and their daily usage become widely adopted by the population, it will be conceivable that the general public might be desensitized to the intimate nature of the data collected. Gaze data is a striking example of how personal preferences and otherwise covert information regarding one’s experience of its environment can be effectively derived. Indeed, metrics such as fixation duration and relative number of saccades when exploring a visual object provides objective measures of how much attention has been paid to the object, gives indication about the depth of cognitive processing of that information, revealing interest and preferences. Advances in the fields of computer vision are providing efficient ways to label visual objects in video recordings in an automatic and increasingly reliable manner. The extraction of neural markers in relation to labeled visual data adds another layer of insight about an individual attention, semantic processing, and decision-making. The fusion of gaze and brain data offer novel opportunities to extend BCI applications beyond the presentation of artificial interfaces and paradigms. It is through the acquisition of such contextual information that neural data will deliver its full potential for real-world applications. Aside of the exciting prospects offered by the exploitation of multimodal data for everyday-life applications, it is critical to reflect on the sensitive nature of such data. Indeed, once equipped with sensors that will become evermore transparent, data will be acquired continuously without the user being conscious of it. The covert aspects of continuous physiological data recordings poses questions regarding the consent of the user for such information to be exploited at any given moment. To be clear, multimodal brain and body imaging data pose novel ethical issues regarding data privacy. It falls upon researchers the responsibility to ensure that data privacy remains a priority by setting precedents of high standards in how such multimodal data is handled.

## 5 CONCLUSION

The challenges related to the study of human cognition in everyday life contexts are numerous. Abandoning computerized paradigm lessens experimental control over a wide range of variables that can add variance to participants’ behaviour and cognitive processes involved in a certain task. In the context of EEG analyses this high degree of freedom poses several conceptual and technical issues, notably related to the timing of experimental and cognitive events. The present findings provide evidence that such challenges may be overcome through a combination of scalable experimental design, recording of multimodal brain and body imaging data, and the application of state of the art signal processing and template matching methods. By applying these approaches, neural markers of cognitive processes related to visual information embedded in a real-world environment could be captured. These encouraging results further highlight the relevance of scalable experiments to study human cognition in real-world contexts. Indeed, recording neural responses elicited by the discrete presentation of visual stimuli through a computerized paradigm (i.e., the tablet condition) was instrumental to create a template that could be used to search for and extract similar responses in a more naturalistic, and therefore less controlled recording setting such as the library condition. Moreover, the extended range of analyses enabled by the joint recording of gaze and brain dynamics showcases the complementarity of mobile brain and body imaging methods. The eye-tracking data provided contextual information regarding the occurrence of experimental events which enabled time-domain analyses to be performed on the EEG data. While the information provided about experimental event timing may not have the temporal resolution required to perform time-domain analyses at the milliseconds scale, they can however serve as an initial estimate upon which template matching approaches are applied to extract ERP features. In conclusion, adopting a scalable approach to experimental design and leveraging the potential of multimodal recording methodologies are important steps toward enabling the study of embodied aspects of human cognition in naturalistic environments.

## Abbreviations

EEG: electroencephalography
E-T: Eye-Tracking
ERP: Event-Related Potentials
MoBI: Mobile Brain/Body Imaging
DTW: Dynamic Time Warping

## Acknowledgements

The authors thank Catriona Bruce and Stephen Stewart for the technical assistance in setting up the experiment. Thank you to Ludovic Darmet for his advice on data analysis.

## Data and code availability statement

Original EEG and E-T data with experimental events annotations will be made publicly available on online repository. The code used for the processing and analysis of the data is also accessible online.

